# ^14^C-age of carbon used to grow fine roots reflects tree carbon status

**DOI:** 10.1101/2024.10.15.618388

**Authors:** Boaz Hilman, Emily F. Solly, Frank Hagedorn, Iris Kuhlman, David Herrera-Ramírez, Susan Trumbore

## Abstract

- “Bomb” ^14^C ages in trees indicate the time elapsed between carbon fixation into nonstructural carbon (NSC) and its use for metabolism and growth. It remains unknown why newly grown aboveground tissues have a narrow range of young ^14^C-ages, while fine root ages can span decades.
- We measured ^14^C in tissues of two coniferous species along an alpine treeline ecotone and used a mixing model to estimate the fraction of NSC that is metabolically active. We expected that greater growth limitation higher in the ecotone would supply more freshly fixed NSC to respiration, active NSC, carbohydrates, and roots, resulting in younger ^14^C-ages of NSC used to grow tissues.
- Results confirmed an increase in fresh NSC supply with elevation. Needle and branch growth was supported by young NSC (< 2 yr) with little elevation influence, while root growth was supported by older NSC that declined from 10 to 6 yr with elevation.
- Massive allocation of fresh NSC to needles and branches could explain their young ^14^C-ages. Variable fine root ^14^C-ages reflect tree C status, with older ages when aboveground growth limits the contribution of new C belowground, while young ages represent greater delivery of fresh NSC to roots.

## Introduction

Of all the processes involved in the allocation of carbon (C) in trees, the flow of C to roots is perhaps the most puzzling. Roots anchor trees to the ground and provide access to soil resources, functions that consume 25—63% of forest C assimilation (Litton *et al*., 2007). To allocate photo-assimilates trees use nonstructural C (NSC). Soluble NSC like sugars is used to transport and supply C to sinks such as respiration and growth. Stored sugars and insoluble lipids and starch represent NSC that is remobilized to survive periods of low photosynthesis (Dietze *et al*., 2014). In the absence of a method to directly measure the flow of phloem sugars, belowground NSC allocation is studied with indirect approaches like measurements of belowground C sinks and isotopic labeling. Under conditions where freshly fixed C exceeds growth demand (“surplus C”), more NSC is typically allocated to nonstructural carbohydrates (NSCarb), respiration, and belowground sinks (Prescott *et al*., 2020). Such conditions have been observed when aboveground growth is limited by non-C resources like nitrogen and water (Marshall *et al*., 2023; Rog *et al*., 2024). Isotopic labeling studies show that during summer fresh NSC is allocated from needles to roots within days, but it takes years to turn over stored NSC stocks, especially in roots (Keel *et al*., 2006; Keel *et al*., 2007; Epron *et al*., 2012; Solly *et al*., 2023). The isotopic composition of NSC pools therefore contains information about allocation processes integrating timescales of years.

A global labeling approach with a resolution of years takes advantage of “bomb” ^14^CO_2_ in the atmosphere, excess radiocarbon produced by nuclear weapons testing in the 1960s. Since then, CO_2_ fixed photosynthetically in fresh NSC each year has a unique ^14^C signature that allows estimation of how long C persists in NSC and tissues (^14^C-age). The ^14^C-age of NSC used by trees to respire and to grow new leaves and wood indicates that carbon fixed within the past few years fuel these C sinks (Harkness *et al*., 1986; McNeely, 1994; Carbone *et al*., 2011; Muhr *et al*., 2013; Andreu-Hayles *et al*., 2015). In contrast to this narrow age span, ^14^C-ages of fine roots (≤ 2 mm) in trees are highly variable and span from < 1 yr to a decade or more (Gaudinski *et al*., 2001; Trumbore *et al*., 2006; Vargas *et al*., 2009; Gaudinski *et al*., 2010). These results were used to suggest slow (many years to decades) fine root turnover, in strong disagreement with multiple observations using minirhizotrons and ingrowth cores documenting a much more dynamic fine root pool (Guo *et al*., 2008; Strand *et al*., 2008). More recent studies targeted this discrepancy by comparing ^14^C ages to root chronological ages, estimated by counting annual growth rings or by collecting tree roots growing through buried screens (Sah *et al*., 2011; Helmisaari *et al*., 2015; Solly *et al*., 2018). The finding of roots with older ^14^C-ages than chronological ages lead to the conclusion that old NSC must be fueling these roots’ growth (Solly *et al*., 2018). However, it is yet unclear why NSC pools in roots are so variable in C ages.

A useful way to understand ^14^C observations is to conceptually separate NSC pools into two functional sub-pools, active and stored [Fig. 1, (Carbone *et al*., 2013)]. We define the active pool as the recently imported NSC that is used to fuel C sinks and typically, but not always, contains fresh NSC. The stored pool has a smaller contribution to metabolism and generally older ^14^C-ages (Herrera-Ramirez, *et al*., 2020). This conceptual separation can explain the younger ^14^C-ages in respired CO_2_ compared to bulk extracted NSC (Hilman *et al*., 2021). It can also explain why older ^14^C is used for respiration and growth when fresh NSC supply to tissues is interrupted (Vargas *et al*., 2009; Carbone *et al*., 2013; Muhr *et al*., 2018; D’Andrea *et al*., 2019; Hilman *et al*., 2021). In the absence of fresh NSC inputs, and assuming ‘last in, first out’ NSC dynamics where the most recent NSC is the most accessible (Lacointe *et al*., 1993; Carbone *et al*., 2013), over time C sinks will access stored NSC with increasingly older ^14^C-ages. Such observations support a common assumption that older ^14^C-ages used in respiration or growth must reflect a greater use of stored NSC.

**Figure 1.**
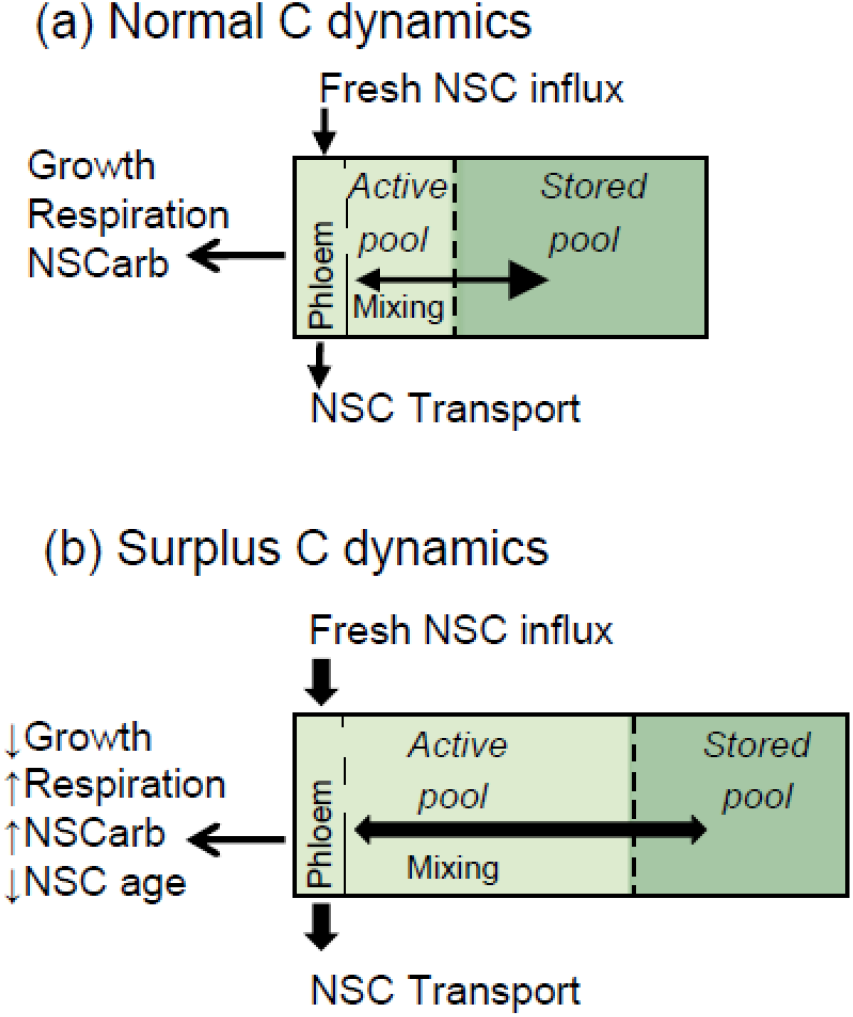
Conceptual model of nonstructural carbon (NSC) pool adapted from Carbone *et al*. (2013), in (a) ‘normal’ and (b) ‘surplus’ C dynamics. Surplus C dynamics frequently occur when growth is suppressed and freshly-fixed NSC is in excess. The boxes and arrow sizes are roughly proportional to the sizes of the pools and fluxes. Freshly fixed NSC enters the active pool via the phloem. The active pool fuels growth, respiration, and stored NSC compounds including nonstructural carbohydrates (NSCarb). Stored NSC can be mixed back into the active pool. The ^14^C-age of the NSC pool is the weighted age of the active and stored pools.

Physically, the two-pool separation can be linked to spatial location in multiannual organs such as roots, with an active region near the cambium and a storage region in xylem tissues, or to molecules’ accessibility (e.g. soluble vs. insoluble forms, or location in cytoplasm vs. in vacuoles and starch granules). ^14^C measurements of individual sugars and starch, however, have been proven to be analytically difficult (Richardson *et al*., 2013; Trumbore *et al*., 2015), therefore researchers rely on operational fractions such as use in respiration or solubility in water to measure NSC, and these inevitably mix active and stored NSC pools. Using a ^14^C mixing model based on comparing ^14^C of extracted NSC with structural tissue to estimate the relative sizes of active and stored fractions, we found that in Eurasian aspen roots the active fraction represented 30% of the total soluble NSC (Hilman *et al*., 2021). In a different study, larger active fractions were suggested to be associated with greater fresh NSC fluxes, higher NSCarb concentrations, and faster metabolism, resulting in faster NSC turnover and younger NSC ^14^C-ages [Fig. 1 (Carbone *et al*., 2013)].

In this study, we tested whether changing allocation of fresh NSC belowground controls root ^14^C-ages. For that purpose we analyzed needles, branches, and fine roots of two coniferous species (deciduous *Larix decidua* and evergreen *Pinus mugo spp. uncinata* Ramond) along the ‘Stillberg’ alpine treeline ecotone. Alpine treeline ecotones are perhaps the most significant example of the influence of growth limitation on surplus C in mature trees (Körner, 1998). At temperatures below 6 °C that prevail at treelines, tree growth is suppressed while C assimilation is maintained (Körner, 2003b; Körner, 2003a). As a consequence of the excess of fresh NSC supply compared to growth demand, trees at higher and colder elevations have a larger surplus C flux, expressed by higher concentrations of NSCarb (Hoch & Körner, 2012). Increase in surplus C at low temperatures is also evidenced by greater root exudation, C allocation to mycorrhiza, and root respiration (Hagedorn *et al*., 2010; Streit *et al*., 2014; Karst *et al*., 2016; Solly *et al*., 2017). To assess surplus C status, we measured respiration rates and NSCarb concentrations (sugars and starch), in addition to ^14^C-ages in respired CO_2_, water-soluble NSC, and structural C. To estimate the age of the NSC used to grow tissues, we compared structural ^14^C-age to roots’ known chronological ages. We also applied the ^14^C mixing model of Hilman *et al*. (2021) to estimate the fraction of the active sub-pool in the soluble NSC. *Larix* is a deciduous and fast-growing pioneer species with higher photosynthesis rates and NSCarb concentrations than Pinus, which has determinate shoot growth (Handa *et al*., 2005; Hoch & Körner, 2009; Barbeito *et al*., 2012; Streit *et al*., 2014). We therefore hypothesized the *Larix* to have faster NSC turnover and younger NSC. We also hypothesized that at higher elevations, greater surplus C would be reflected by a larger NSC active fraction and younger ^14^C ages (Fig. 1).

## Materials and Methods

### Study site and experimental design

The ‘Stillberg’ site (Swiss Alps, 46°47⍰N, 9°52⍰E) hosts an afforestation experiment established in 1975, where *Larix* and *Pinus* trees (and *P. cembra* that died out) were planted in equal distribution and abundance (Barbeito *et al*., 2012). The uniform ages allowed us to avoid potential effects of tree age on ^14^C (Vargas *et al*., 2009) and focus on effects of elevation on C allocation. Over 200 m of elevation we selected three “ecotone” sites at 2000 (“low”, outside the afforestation, trees with the same age), 2080 (“middle”), and 2200 (“high”) m a.s.l. For an additional comparison we sampled trees planted in the same year from a “valley” site located at 1600 m a.s.l., where temperatures are higher, and trees are 2–6 times taller than the ecotone trees. At the ecotone, slowing growth rates with elevation result in tree height decline, from 10 m (low) to 3 m (high) for the *Larix* and from 2 m (low) to 1.5 m (high) for the Pinus. Average air and soil temperatures during the growing season decrease with elevation from low to mid-elevation, but at high elevation, while air temperatures are still decreasing, soil temperatures increase and are warmer than the air (air 8.4 °C, soil 8.9—9.4 °C), probably because of more direct sunlight related to widespread tree mortality and shorter tree stature (Hilman *et al*. unpublished). In spite of the higher average temperatures, the time of snowmelt at the high elevation is at least two weeks later than at the lower elevations.

Over three days in the first week of September 2019 we harvested tissues from five replicate trees per species and elevation in the ecotone sites, and from three replicate trees per species in the valley. To avoid isotopic contamination, trees were at least 50 m distant from sites of previous studies that used free air CO_2_ fertilization (FACE) and ^14^C labeling in pots (Dawes *et al*., 2015; Ferrari *et al*., 2016). From each tree we collected 2-yr old branches including needles, and roots. To be sure the roots were intact and belonged to the studied tree we tracked them back to the base of their tree stem. In the evening of every harvest day we performed respiration incubations of branches and water-washed fine-root clusters. Incubated roots were dried and used to measure root chronological ages. In parallel, we prepared a second set of samples for NSC extraction by placing them in a microwave (2 min on 900 W) to halt carbohydrate consumption processes. Later we oven-dried the samples at 60 °C for at least two days. We sorted the dry roots by diameter using a caliper to separate them into three size classes: < 0.5 mm, 0.5–1 mm, and 1–2 mm. The needles were removed entirely from the branches, and then the branch bark was scraped off to focus on the wood and be consistent with previous treeline studies (Hoch & Körner, 2012). For NSC extractions samples were milled and homogenized using a ball-mill, except for the finest roots (< 0.5 mm) that contained only a small amount of material. To avoid loss of material during milling of these roots, we used a blade to cut them into small pieces for analysis. In the same field campaign, we buried in-growth screens across the ecotone. The screens were made from 1-mm nylon mesh glued to PVC collars (Ø 25cm, width 5 cm). In September 2020 we collected roots growing through the screens. Identification of the origin of the roots was difficult because many roots were torn during excavation, but un-identified roots were analyzed only if they were suberized.

### Respiration measurements

We incubated fresh whole root clusters (≤ 2 mm) and barked branches for one day and collected the respired CO_2_ in glass flasks for later analysis. One day was the time assumed necessary to collect enough CO_2_ for ^14^C analysis. The incubation set-up included gas-tight Plexiglas cylinders connected on each side with one flask equipped with a LouwersTM O-ring high-vacuum valve (LouwersHanique, Hapert, Netherlands) (Muhr *et al*., 2018). The total headspace was 270–450 cm^3^. Flasks were prefilled with CO_2_-free air (20% O_2_, 80% N_2_), but the cylinder headspace contained local air with ambient ^14^CO_2_. To account for the extraneous CO_2_ we sampled the local air with duplicate 2 L flasks on each measurement day. For ^14^C measurements we converted CO_2_ to graphite using iron catalyzed reduction with H_2_ (Steinhof *et al*., 2017). Graphitized samples were analyzed by accelerator mass spectrometry (AMS; Micadas, Ionplus, Switzerland) in the radiocarbon laboratory in Jena, Germany. Radiocarbon data are expressed as Δ^14^C, the deviation in permil of the ^14^C/^12^C ratio from “Modern” C (Trumbore *et al*., 2016):

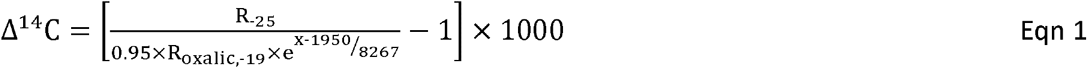

Where R_-25_ is the ^14^C/^12^C ratio of the sample corrected for mass dependent fractionation by normalizing the sample’s δ^13^C to a δ^13^C of −25‰. R_oxalic,-19_ is the ^14^C/^12^C ratio in the standard, oxalic acid, normalized to δ^13^C of −19‰, and the 0.95 term converts to the absolute radiocarbon standard (1890 wood) activity in 1950. The exponent corrects for decay of ^14^C in the standard between 1950 and the year of measurement (x) using mean-life of 8267 yr, to provide the absolute amount of ^14^C in our samples.

To estimate CO_2_ efflux rate we measured the CO_2_ concentration in one duplicate flask using an isotope ratio mass spectrometer (Delta+ XL; Thermo Fisher Scientific) coupled to a modified gas bench with Conflow III and GC (Thermo Fisher Scientific, Bremen, Germany). The mean CO_2_ efflux during the incubation period (mg C g^-1^ day^-1^) was calculated using:

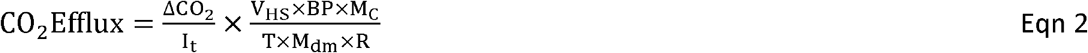

where ΔCO_2_ is the net change in CO_2_ concentration during the incubation (ppm/10^6^), I is the incubation time (days), V_HS_ is the volume of the headspace (mL), BP is the local barometric pressure (hPa), M_C_ is the molar mass of C (12 mg mmol^-1^), T is the temperature of incubation (K), M_dm_ is the dry mass of sample (g), and R is the ideal gas constant (83.14 mL hPa K^-1^ mmol^-1^).

Respiration rates are related to temperature. Incubation experiments were done in a stone barn with air temperature of 10.1 ± 0.6 °C (mean soil temperature across all ecotone sites was 11.1 ± 2.1 °C) and are reported without further correction. Data are not reported for several samples from the valley and low sites, however, as these incubations were exposed for several hours to higher temperatures after leaving the site.

### Extractions of nonstructural and structural carbon

We extracted samples sequentially for water-soluble NSC (including soluble sugars), starch, and α-cellulose. The water-soluble C was extracted by mixing of 50-mg samples in 1.5 ml deionized water at 65 °C (Landhäusser *et al*., 2018; Hilman *et al*., 2021) in Eppendorf tubes that were extensively prewashed to leach any potential extraneous C contribution. A sub-sample from the eluate was used to quantify the concentrations of the sugars fructose, glucose, and sucrose using high-performance anion-exchange chromatography with pulsed amperometric detection device (HPLC-PAD, Dionex® ICS 3000, Thermo Fisher GmbH, Idstein, Germany) (Raessler *et al*., 2010). We express sugar concentrations in units of glucose equivalent per dry mass. The rest of the eluate was oven-dried (40 °C), graphitized, and analyzed in the AMS for ^14^C.

Starch from the water-extraction pellet was digested by two enzymes (Landhäusser *et al*., 2018): α-amylase (Sigma cat. no. A4551) that converts the starch to water soluble glucans, and amyloglucosidase (Sigma cat. no. ROAMYGLL) that converts the glucans to glucose, which was measured in the HPLC. As for the sugars we express starch concentrations in glucose equivalent (Landhäusser *et al*., 2018).

α-cellulose from the remaining pellet was extracted and measured for Δ^14^C (Steinhof *et al*., 2017). α-cellulose is immobile and thus commonly used as a proxy for structural biomass (Leavitt & Danzer, 1993). The extraction is based on toluene-ethanol removal of lipids, bleaching by sodium chlorite and acetic acid to remove lignin, and isolation of the cellulose by strong base (sodium hydroxide). The α-cellulose is then graphitized and analyzed for ^14^C.

### ^14^C-age estimates

Estimation of bomb ^14^C-age is based on comparison between the ^14^C signature of a sample and the local record of atmospheric ^14^CO_2_. We recently estimated the atmospheric record in Stillberg (Hilman *et al*. unpublished) according to data from local tree-stem rings, direct atmospheric Δ^14^CO_2_ measurements in two near stations [Jungfraujoch, Swiss alps and Schauinsland, the Black Forest, Germany, (Hammer & Levin, 2017)], and a global compilation of ^14^C data from different sources for the region (Hua *et al*., 2013). The local atmospheric Δ^14^CO decreased linearly since 2000 at a rate averaging 4.5‰ per year to a value of −0.9‰ in 2019 when most of the samples were collected, and −5.4‰ in 2020 when we collected the screen roots. For samples with Δ^14^C values within the range of the years 2000-2020 we calculated the mean C age with the equation:

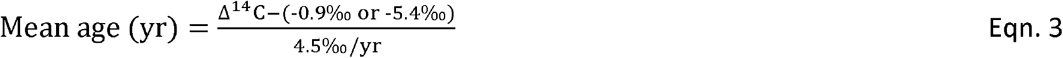

For samples with Δ^14^C values higher than the 2000 atmospheric Δ^14^CO_2_, we estimated the age by matching the Δ^14^C to the year with the closest value in the Hua *et al*. (2013) dataset.

### Chronological ages

Chronological ages represent the lifespan of a tissue. For tree branches it is possible to determine the chronological age from the branch position, corresponding to 2 years for all of our samples. The evergreen *Pinus* needles thus have ages of 0–2 years. The chronological age of the needles of the *Larix* was assumed to be 0 yr because they are deciduous. We estimated fine-root chronological ages according to Solly *et al*. (2018). In brief, *Larix* and *Pinus* present discernible boundaries between two neighboring growth increments (Cutler *et al*., 1987; Mrak & Gričar, 2016), which can be used to count annual growth rings in fine roots (Solly *et al*., 2018). We counted the growth rings in fine roots for all samples by randomly choosing at least three individual root segments. Some samples did not have sufficient roots for all replicates in some of the size classes. Cross thin sections (approximately 15–20 μm) of the fine roots were prepared using a laboratory microtome (Gärtner & Schweingruber, 2013), and subsequently covered with glycerol and a cover glass. The thin sections were photographed with a BX41 system microscope (Olympus, Tokyo, Japan). The number of growth rings was counted in the secondary xylem of the root thin sections. Only complete growth rings were taken into account during the measurements (Solly *et al*., 2018). In case no growth rings were present, or a secondary xylem was absent, the roots were considered to be younger than one growing season (that is, 0.5 yr). We assumed a chronological age of 0 yr for the roots sampled through the root screens.

### Estimates of the age of NSC used for growth

We assume that the α-cellulose ^14^C-age is the sum of two components (Gaudinski *et al*., 2001): the time between C fixation and tissue synthesis (when the C resides in NSC), and the time between the synthesis and the year of sampling (equal to chronological age). Accordingly, the age of the NSC used to grow new tissues was determined by subtracting the chronological age from the α-cellulose ^14^C-age. The chronological age, however, represents the number of years since first tissues began to grow in the organ (e.g. inner growth ring), but tissues that grew in later years (e.g. external growth rings) have younger chronological ages. The chronological age is therefore not comparable with ^14^C-age that integrate across all tissue growth years, and for samples with chronological age > 1 yr we used the mean chronological age, i.e. slightly younger than the chronological age.

### Statistical analysis and mixing model

To test effects of elevation, species, and tissue we applied linear mixed-effects models with the “lmer” function in the R package “lme4” (Bates *et al*., 2015; R Development Core Team, 2019). Starting from a null model with individual tree as random intercept, we added sequentially the fixed factors and their two-way interactions. The increasingly complex seven models were tested for parsimony – that is whether the increased complexity improves significantly the model ability to explain the data. Parsimony was tested using the “anova” function where a χ^2^ test was applied to assess the statistical significance of stepwise model improvement. When factor or interaction was significant, we applied estimated marginal means (EMMs) post-hoc test to compare factor levels [using “emmeans” function in the package “emmeans” (Lenth, 2021)]. The response variable was square root (SQRT) or log transformed when the assumption of homogeneity of variance was violated. For NSCarb comparisons, which varied greatly among organs, we standardized concentrations by dividing concentration of each sample with the mean concentration of the respective tissue type and species (Hoch & Körner, 2012).

We used a mixing model to estimate the fraction of active NSC from the total water-soluble NSC. The model assumes that the ^14^C content of structural C is correlated with the stored NSC but not with the active NSC (Hilman *et al*., 2021). The former is explained by concurrent allocation of NSC to the stored NSC and structural C. If 100% of the soluble NSC is “stored”, a linear regression between soluble NSC and structural C in different roots will have a slope of 1. Existence of an active pool with recently translocated NSC (fresh or from storage in other tissue), with a ^14^C signature not yet passed into structural C, will alter the slope of the regression. We calculated the regression slope with the lmer function for the relation soluble NSC age ∼ α-cellulose age for different tissues and for roots from different elevations. The random effect was the species when we compared tissues and individual tree when roots in different elevations were compared. To extract the slope estimates and their P values we used the function “summary” from the “lmerTest” package (Kuznetsova *et al*., 2017).

## Results

The species difference was not significant for most of the measured variables. For clarity of presentation, species have been pooled in most figures.

### Nonstructural carbohydrates and respiration rates increase with elevation

Standardized concentrations of total nonstructural carbohydrates (NSCarb, soluble sugars + starch) increased by 27% from the bottom to the top of the ecotone (Fig. 2a; χ^2^ = 8.54, P < 0.04). The elevation increase was mainly driven by the sugars (χ^2^ = 11.10, P < 0.02) while starch was constant across elevations (χ^2^ = 5.68, P = 0.13). We detected a stronger statistical significance when only fine-root sugars were analyzed (χ^2^ = 12.18, P < 0.007), without needles and branch wood (Fig. 2b). Respiration rates across tissues were also highest at the high elevation (Fig. 2c; χ^2^ = 11.09, P < 0.004). Absolute NSCarb concentrations declined by a factor of five from needles to the finest roots (χ^2^ = 165.90, *P* < 0.001) and were 40% higher in the *Larix* than in *Pinus* (Fig 2d; χ^2^ = 7.00, *P* < 0.009). Of the total NSCarb mass, sugars were the majority (≥ 85%) in needles, branch wood, and < 0.5 mm roots, while starch was the majority (30–50%) in the larger roots

**Figure 2.**
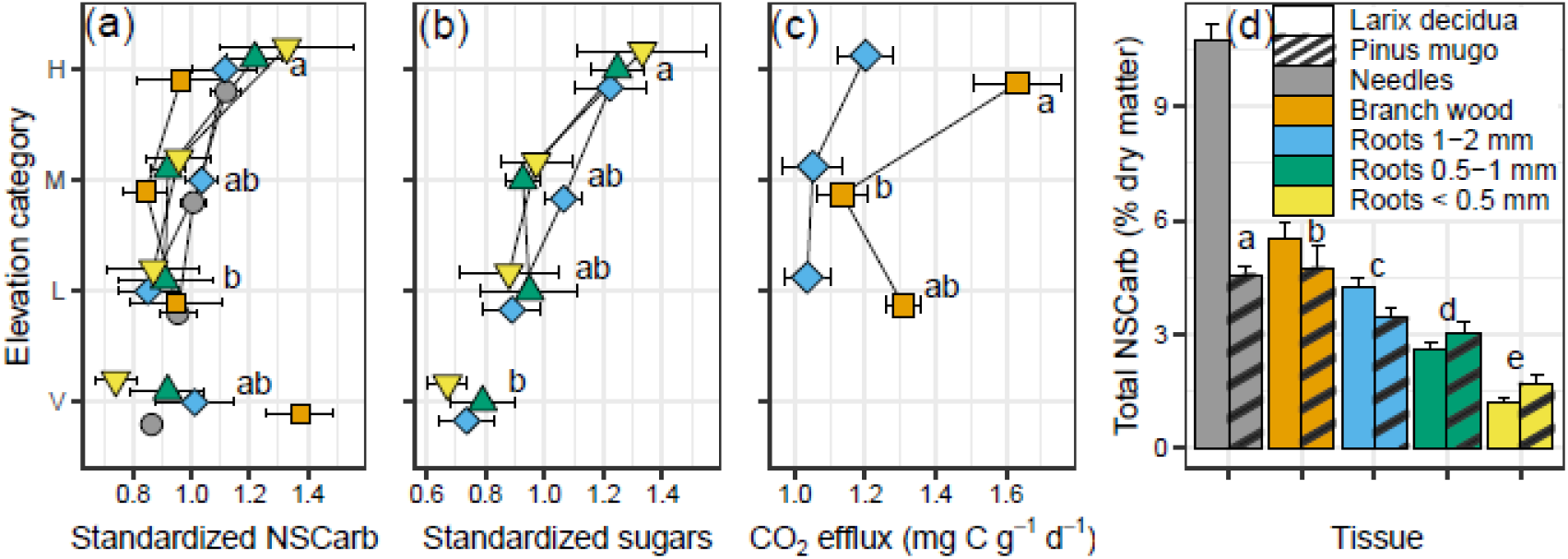
Nonstructural carbohydrates (NSCarb) and respiration rates. (a) Elevation × organ effect on standardized NSCarb (soluble sugars + starch) concentrations (means ± SE; n = 6– 10). Concentration for each tissue type was standardized by dividing its measured value by the mean concentration of the respective tissue and species. H, high elevation; M, middle elevation; L, low elevation; V, valley, not in the ecotone. (b) Standardized soluble sugar concentrations in the roots. (c) Respiration rates (CO_2_ efflux) from excised branches including bark and roots (≤ 2 mm) incubated at 10.1 ± 0.6 °C (n = 4–10) (d) Tissue and species effects on NSCarb concentrations (n = 15–18). Different letters represent significant differences (P < 0.05) between elevations in panels a, b, and c, and between tissues in panel d.

### ^14^C ages and mixing model

The greatest variability in ^14^C-ages was observed among tissues (Fig. 3 and Table S1). The ^14^C-age of α-cellulose of needles (1.3 ± 0.2 yr, mean ± SE) was slightly younger than that of branches (2.7 ± 0.3 yr). The three fine-root size classes were significantly older than the aboveground tissues, but did not differ from each other (1–2 mm: 9.3 ± 0.9 yr, 0.5–1 mm: 8.3 ± 0.9 yr, < 0.5 mm: 8.4 ± 0.9 yr). The soluble NSC ages were usually younger than the α-cellulose with similar tissue effect (Fig. 3a, b and Table S1). In addition, soluble NSC was the only fraction (including respired CO_2_) that differed between species, where *Larix* contained younger NSC than *Pinus* (4.1 ± 0.3 yr vs. 5.8 ± 0.4 yr; χ^2^ = 11.51, P < 0.001). The elevation effect was strongest when interacting with tissues (Table S1): α-cellulose in the aboveground tissues had statistically equal ages across elevations (despite slightly higher branch ages at mid elevations), in contrast to fine roots that varied widely. Averaged across all size classes and species, α-cellulose age of fine roots was youngest at the valley (3.8 ± 0.9 yr) and oldest at the bottom of the ecotone (11.9 ± 0.9 yr). From the bottom to the top of the ecotone the mean age decreased to 8.2 ± 0.7 yr. Fine roots respired slightly older CO_2_ (1.4 ± 0.2 yr) than the branches (Fig. 3c; 0.7 ± 0.2 yr; χ^2^ = 5.45, *P* < 0.02). The ^14^C-ages of the respired CO_2_ correlated with the soluble NSC ages, and the regression equation suggests that about 85% of the respired CO_2_ originates from < 1 yr NSC (intercept of 0.35 yr) and 15% originates from the local NSC, which is older in the roots (Fig. 4).

**Figure 3.**
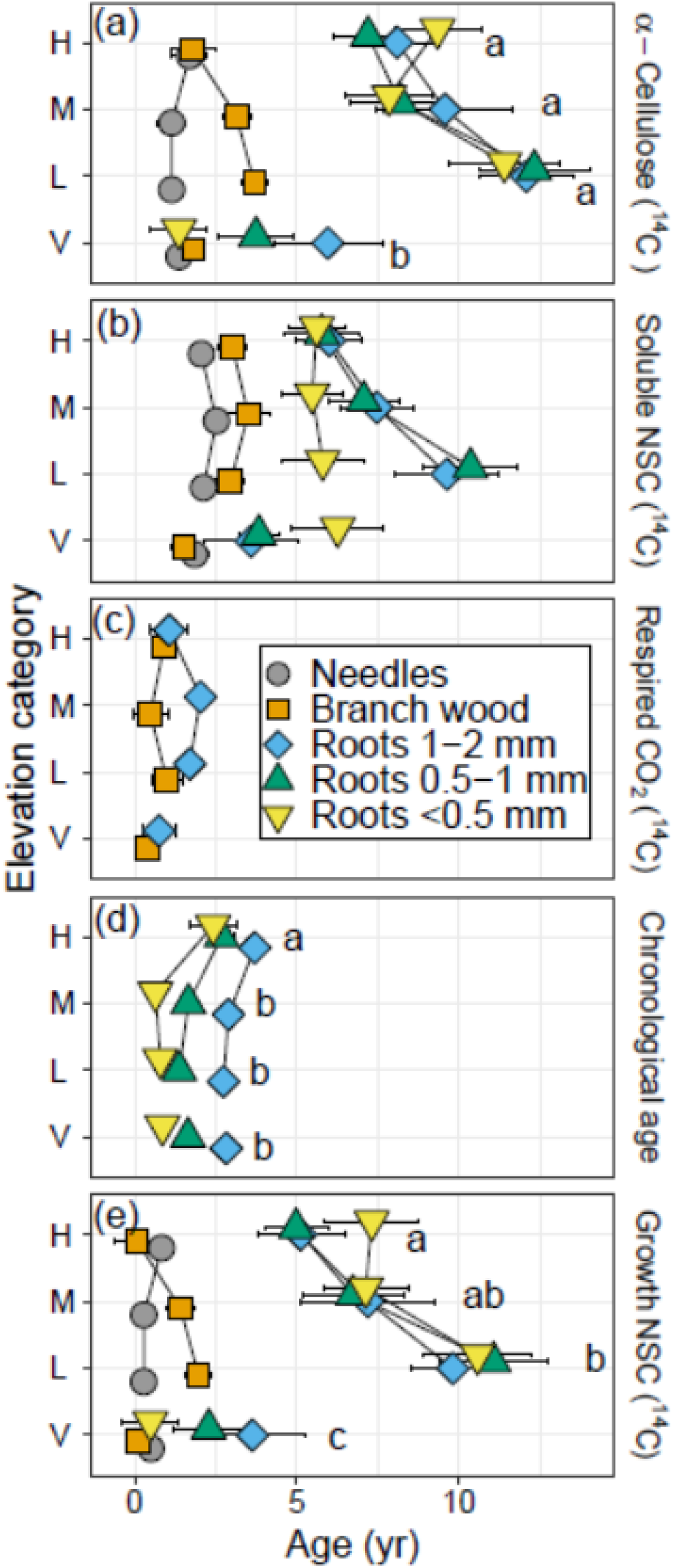
^14^C-ages of carbon pools averaged by elevation × organ (means ± SE; n = 6–10). (a) ^14^C-ages of structural α-cellulose. (b) ^14^C-ages of the nonstructural carbon (NSC) extractable in warm water. (c) ^14^C-ages of CO_2_ respired by incubated fresh branches including bark and roots (≤ 2 mm). (d) Chronological ages estimated by counting annual growth rings in the secondary xylem of fine roots. (e) ^14^C-ages of the NSC used to grow new tissue calculated by the difference between the α-cellulose age and the chronological age, corrected for mass contributions of each annual ring. H, high elevation; M, middle elevation; L, low elevation; V, valley, not in the ecotone. Different letters represent significant differences at *P* < 0.05 between elevations, for the interaction elevation × organ when all root sizes where pooled together.

**Figure 4.**
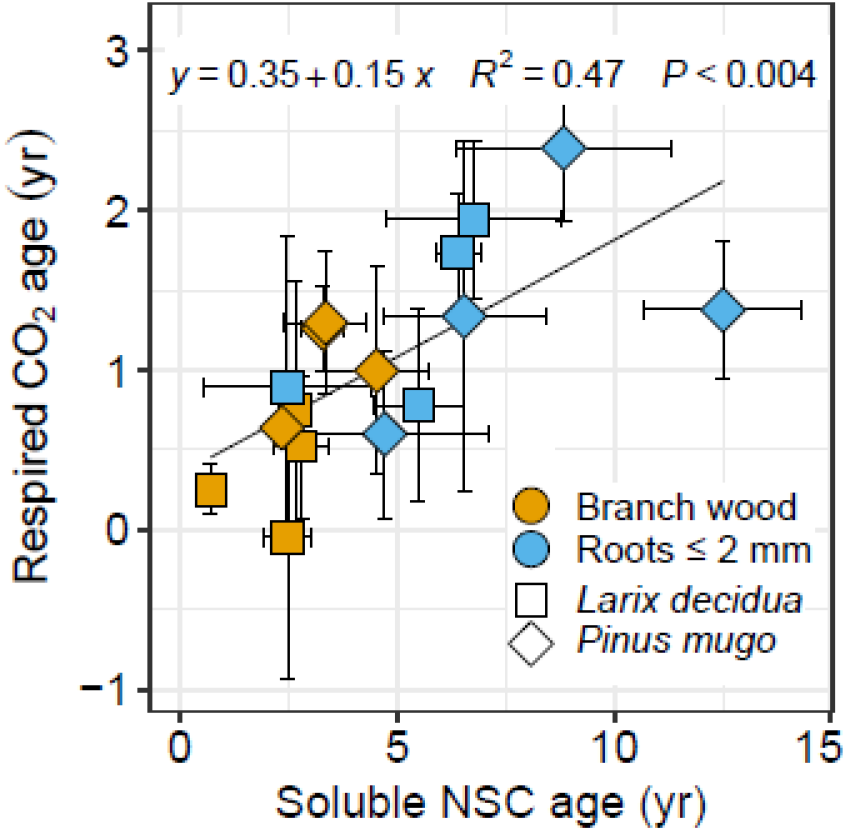
Relationship between the ^14^C-ages of respired CO_2_ and soluble NSC. Each data point is the mean ± SE of each organ × species × elevation combination (n = 3–5). The equation and statistics of a linear regression are presented.

Our mixing model shows that at the bottom of the ecotone, where roots had the oldest ^14^C-ages, the active pool fraction is the smallest (14%, Fig. 5a). The decrease in NSC ages with elevation parallels an increase in the active NSC fraction (44-47% at mid and high elevations). In the valley, where root NSC ages were youngest, the active pool fraction is greatest (84%). The inverse relationship between NSC age and active fraction was also observed among tissues (Fig. 5b), where needles and branch wood with young NSC ages had large active fractions (67-93%), while the two larger root classes with old NSC ages had small active fractions (28-32%). The < 0.5 mm roots stand out as having large active NSC pools and old NSC ages.

**Figure 5.**
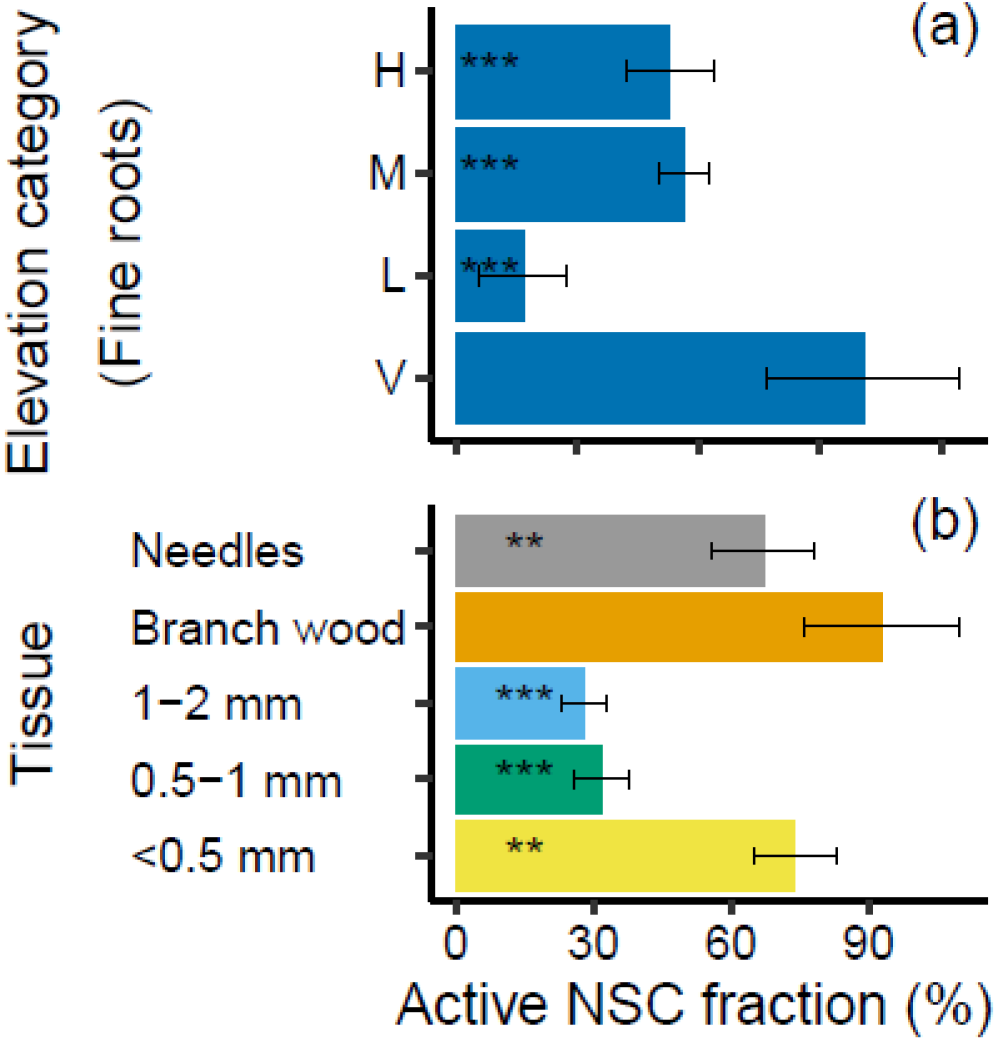
The fraction of the metabolically active pool in the total nonstructural carbon (NSC). (a) Active pool fraction in fine roots (≤ 2 mm) when all root sizes of both species where pooled together. H, high elevation; M, middle elevation; L, low elevation; V, valley, not in the ecotone. (b) Per cent of NSC in the active pool for different tissues when both species where pooled together. Diameter categories refer to fine root classes. Asterisks show the significance of the estimate in the statistical model: ‘***’ <0.001, ‘**’ <0.01, ‘*’ <0.05. Error bars represent the standard error of the model estimate.

### Chronological and growth NSC ages

For the aboveground tissues we estimated ages of 0–2 years according to stem branching and foliar type. Growth-NSC ages (mean chronological age subtracted from α-cellulose ^14^C-age) for all needles and branch wood in the high elevation and valley trees were ≤ 1 yr, while at mid-elevations trees used 1–2 years old NSC to grow branch wood (Fig. 3e). The chronological ages of fine roots ranged between < 1 to 3.6 yr, with older ages in *Pinus* over *Larix* (χ^2^ = 4.29, *P* < 0.04), older ages in thicker roots (Fig. 3d; χ^2^ = 90.73, *P* < 0.001), and oldest ages at the highest elevation (χ^2^ = 13.60, *P* < 0.004). Despite similar chronological ages to the aboveground tissues, fine roots grew from much older NSC (Fig. 3e; tissue effect, χ^2^ = 118.37, *P* < 0.001), in an elevation pattern that tracks the soluble NSC and α-cellulose; oldest growth ages at the bottom of the ecotone (10.5 ± 0.9), younger ages at the top of the ecotone (5.8 ± 0.7), and the youngest at the valley (2.2 ± 0.8; elevation × tissue effect, χ^2^ = 29.54, *P* < 0.001).

The mean growth-NSC age for screen roots was 9.1 yr with no elevation effect, fairly close to 7.8 yr, the mean growth-NSC age estimated for the standing-biomass roots in the ecotone (Fig. 6). We could not validate the origin of 19 woody roots that could not be traced to a specific tree, but the mean ^14^C-age for 6 validated samples from the investigated trees is close to the overall mean, 9.5 yr. While the individual roots collected from screens and the bulked and ground standing biomass roots had similar mean ages, their age distributions differed, with more normal distribution for the standing biomass roots and right skewed distribution for the screen roots with median age of 5.1 yr.

**Figure 6.**
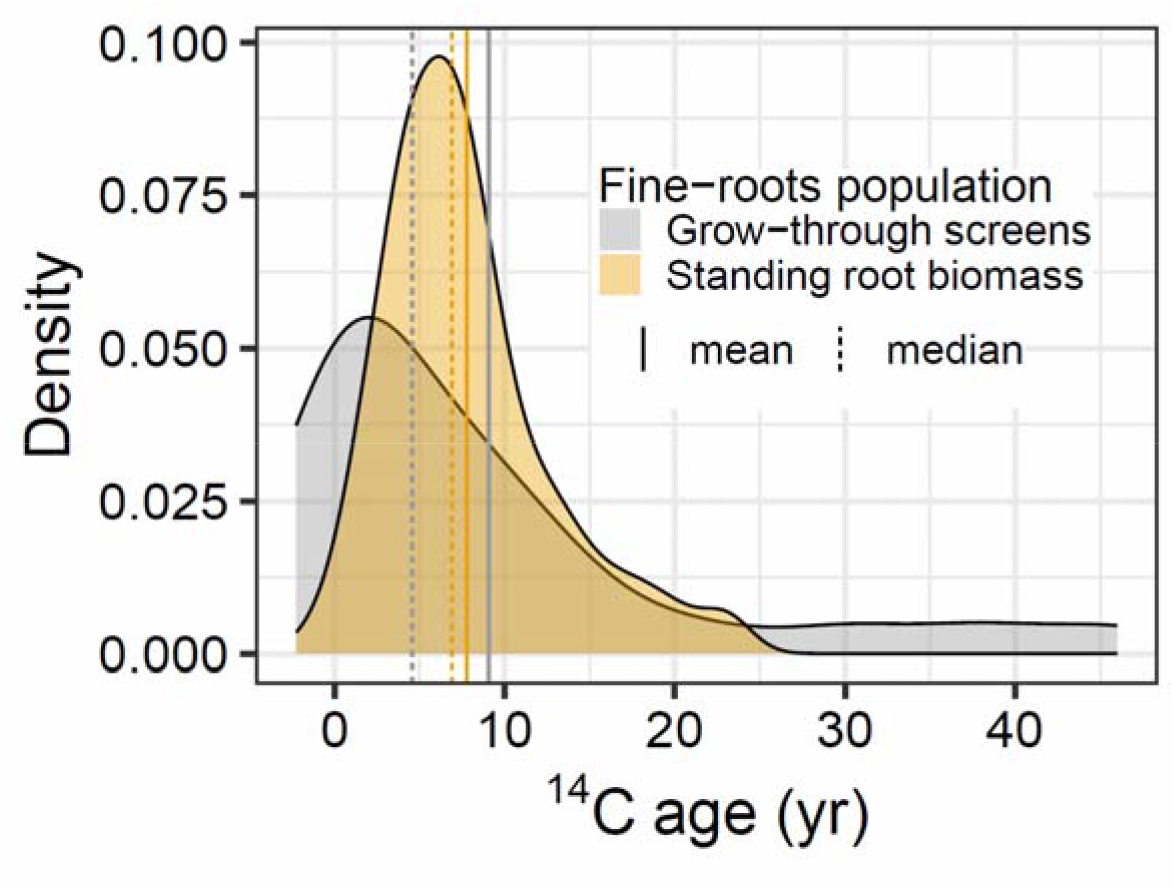
Probability density of ^14^C-ages of NSC used for the growth of fine-roots. The roots are either from roots growing through installed roots screens with known chronological age <1 year (n = 25) or fine roots in standing biomass (i.e. samples in Figure 3e; n = 89).

## Discussion

We suggest that influx of fresh NSC into tissues over years shapes the distribution of ^14^C-ages in tree NSC and structural tissues. In needles and branches the influx (and efflux) is always high, and they are physically closer to the source of fixed carbon, therefore the ^14^C-ages of NSC and tissues are young. Influx of fresh NSC into fine roots, however, is sensitive to metabolism aboveground and in coarse roots; young ^14^C-ages in fine roots reflect a surplus of fresh NSC to the belowground, such as when aboveground growth is limited by temperature more than C assimilation at the high elevation. Older fine root ^14^C-ages at low- and mid-elevations reflect smaller surplus and more efficient use of fresh NSC in needles, stems, and coarse roots, limiting the supply of fresh NSC to the roots. Figure 7 summarizes our results with the proposed fluxes.

**Figure 7.**
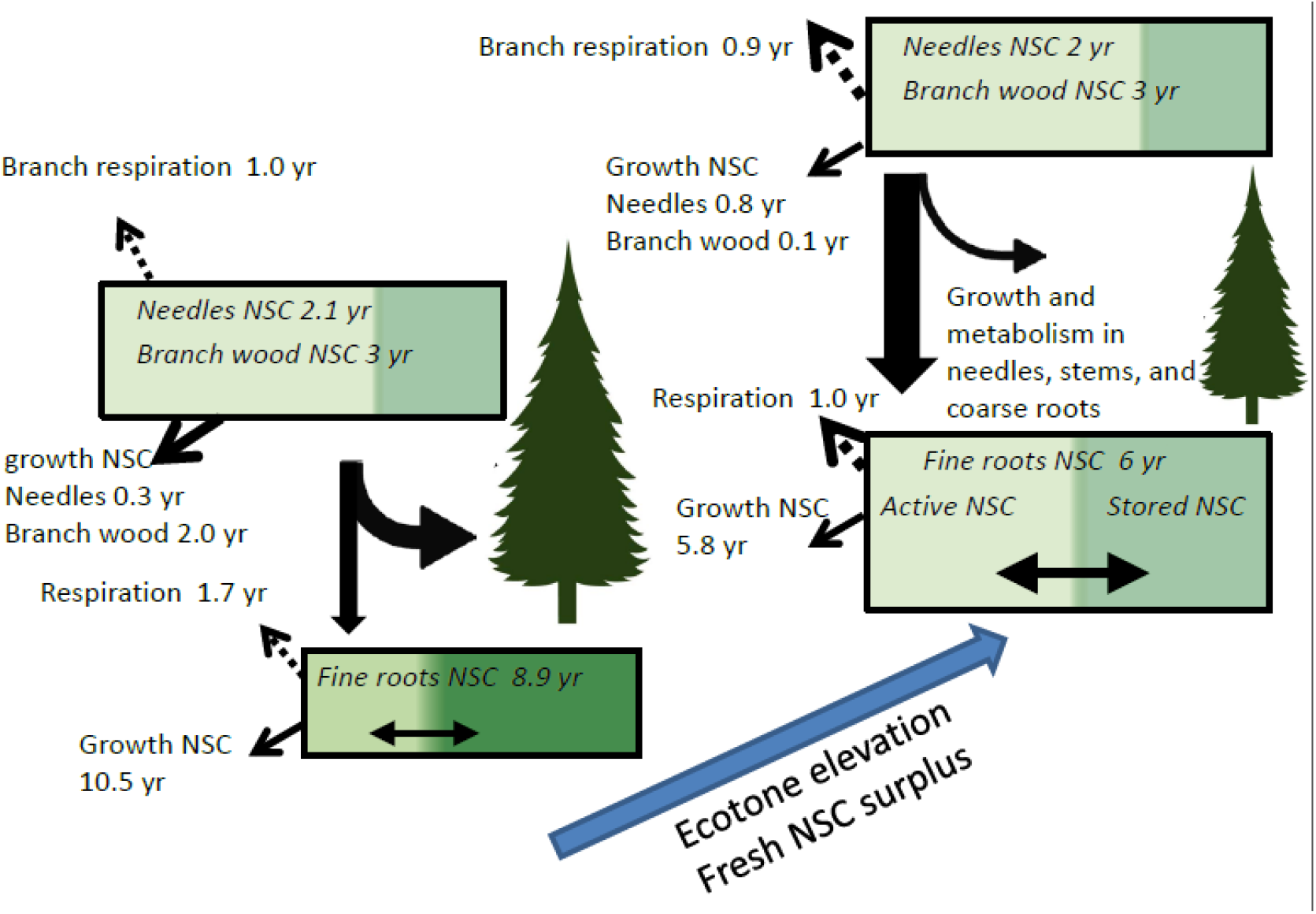
Schematic drawing of the mean ^14^C ages we measured in the bottom and top of the treeline ecotone and suggested fluxes. Box and arrow sizes are roughly proportional to pool and flux sizes. The active pool fuels growth, respiration, and stored NSC compounds, although stored NSC can mixed back into the active pool. The ^14^C age of the NSC pool is the weighted age of the active and stored pools.

### ^14^C ages of fine roots are a sensitive indicator for tree C allocation

We found clear evidence for increased ^14^C-age of NSC with distance from leaves within a tree. Parallel to the increase in needle-to-root age we observed, with exception of the < 0.5 mm roots, a general decrease in the modeled active NSC fraction (Fig. 3b, 5b). The mixing model assumes that the active fraction contains recently transferred NSC, so a larger active fraction implies a larger influx of NSC, which is usually fresh. Accordingly, the young NSC in needles and branches results from a high influx of fresh assimilates. This is consistent with the high metabolic rates of these tissues and with the fact that NSC allocated to the rest of the tree must pass through these tissues. Since fine roots are the most distant tissues from the needles, upstream NSC consumption reduces the amount of fresh sugars reaching fine roots. The < 0.5 mm roots are exceptional by having a large active pool, similar in size to the needles, but with older soluble NSC (Fig. 3b, 5b). A large active NSC pool in the finest roots is not surprising as this size class contains absorptive roots that compared to the coarser transportive roots have faster metabolic activity and NSC turnover (McCormack *et al*., 2015). The old NSC suggests that when fresh C supply is less than demand, multi-year storage is mobilized to the roots.

Our sampling was done in late summer when fresh NSC dominated root respiration (Fig. 4) and C transferred to symbiotic ectomycorrhizal fungi at the site (Hilman et al. unpublished). We can therefore assume that fresh NSC also supported root growth during summer. Pulse-labeling experiments of boreal trees showed a similar allocation of fresh NSC to roots during summer, while earlier in the growing season fresh NSC was mainly allocated to aboveground tissues, rather than to roots (Kagawa et al., 2006; Högberg et al., 2010). We therefore conclude that the measured old NSC used to grow new root biomass (Figs. 3e, 6) represents root growth early in the growing season when, in the absence of fresh NSC, growth is supported by old storage C. The similar NSC used for growth among the three fine root size classes further suggests that they share the same NSC substrate, perhaps exchanged within the larger coarse root system. The shared substrate could also explain why root size and order sometimes have little influence on ^14^C-ages (Gaudinski et al., 2010; Hilman et al., 2021). Thus, the old NSC that fueled growth of the < 0.5 mm roots may also have contributed to the stored NSC pool, and the strong inputs of fresh NSC in the summer were insufficient to turn over the old inherited NSC.

We initially hypothesized that at higher elevations, low air temperatures would limit tree growth but not photosynthetic rates. This would lead to a greater flux of surplus C that would reduce ^14^C-ages of root NSC and α-cellulose. The results support our hypotheses. The historical influence of low temperatures on tree growth was evident from the decrease in tree size with elevation. We additionally found evidence of a more contemporary elevational difference in growth limitation, as indicated by the oldest root chronological ages at high elevation that reflect the longer time it took for a root to grow and reach our diameter thresholds (Fig. 3d). While growing-season soil temperature at high elevation was not colder than at lower elevations (Hilman et al. unpublished), the slower root growth rates can be explained by the shorter growing season at high elevation, indicated by the later snowmelt. During the growing season, the soil at the high elevation was warmer than air, thus the surplus C due to low aboveground C demand could be enhanced by high C demand in the warm roots. Indications for an increase in surplus C and belowground allocation of fresh NSC with elevation are the increases with elevation of the standardized NSCarb, respiration rates, and active-pool fraction in fine roots (Fig. 2a-c, 5b). The carbohydrates that controlled the NSCarb trend are soluble sugars that are known to both improve cold tolerance in plants (Tarkowski & Van den Ende, 2015) and promote root exudation (Farrar *et al*., 2003; Karst *et al*., 2016). The ^14^C-ages in the fine roots also follow the expectation, with an age decrease from the bottom to the top of the ecotone (Fig. 3a,b,e). Aboveground tissues had small ^14^C variability across elevations and contained mainly young NSC, except the branch wood that grew from slightly old NSC (up to 2 yr) at mid elevations. At these elevations trees were the largest (among ecotone trees) and fine root ages were the oldest, suggesting that old ^14^C-ages are linked with smaller surplus C and tighter C balance with more efficient conversion of photo-assimilates into aboveground biomass.

The co-variation of the two species across elevations, despite their different life strategies, implies that tree species might be less important than environmental conditions in controlling ^14^C-ages and NSC allocation. The *Larix* is a deciduous tree with higher growth, photosynthesis, and NSCarb concentrations than Pinus. We therefore expected faster NSC turnover in the *Larix* with younger ^14^C-ages. The soluble NSC of the *Larix* was indeed younger, but only by ∼ 2 yr, and the ages of the NSC supporting respiration and growth were statistically equal among species. The overall similar ^14^C-ages of the two species indicate that the higher *Larix* NSCarb concentrations reflect a large fraction of carbohydrates with fast turnover that does not mix into the multi-year NSC stocks. Among elevations, the youngest root ages were found in the valley site below the ecotone, from which we would infer the greatest allocation of fresh NSC belowground. However, the growth of these trees was the least limited as apparent by their taller stature. Other factors that influence NSC allocation and could explain the young ages in the valley trees are nutrient or water availability, as observed in irrigation of a dry Swiss forest that increased belowground allocation and halved the ^14^C-ages of fine roots (Herzog *et al*., 2014; Gao *et al*., 2021). The suggestion of a tight link between C allocation and ^14^C-ages in fine roots is exciting and may provide new means to investigate the difficult yet important flux of C directed to roots. We suggest that future studies would aim to link more quantitatively between C allocation and ^14^C-ages, by detailed experiments and models for example. Those links could provide spatial and global trends of tree C status and belowground allocation from ^14^C measurements.

### ^14^C-age variability does not reflect necessarily variation in storage use

Older ^14^C-ages in C sinks indicate storage time in the tree, but not necessarily in the tissue. As we concluded above, C allocation dynamics can vary the ^14^C-ages of stored NSC among tissues and sites. This understanding has implications for how ^14^C measurements are used to infer storage use. Here, we measured older CO_2_ respired by fine roots compared to branches (Fig. 3c). While one can conclude that the roots used a greater fraction of stored NSC for respiration, the correlation between the respired CO_2_ and soluble NSC ages (Fig. 4) suggests a constant 15% of the respired CO_2_ originates from local stored NSC, which is older in the roots. To estimate the fraction of storage use in respiration information about the local NSC is required. This can be assessed by the same correlation approach as applied here, or by repeated incubations of excised tissues (Hilman *et al*., 2021; Peltier *et al*., 2023). In this approach, tissues are forced to respire storage NSC and the ^14^C of the stored fraction can be estimated.

### Isotopic proxies are inadequate to estimate fine root lifespan

The old NSC used to grow new fine roots confirms previous observations of the inconsistency between root longevity and ^14^C-ages (Solly *et al*., 2018). Our fine-root chronological ages are in line with fine-root longevity of months to ∼3 yr estimated by minirhizotrons and sequential coring for coniferous trees growing in similar environmental conditions (Guo *et al*., 2008; Brunner *et al*., 2013). Non-isotopic approaches provide no evidence for widespread fine-root ages of decade or two as suggested by the early ^14^C studies (Gaudinski *et al*., 2001; Riley *et al*., 2009; Gaudinski *et al*., 2010). Besides ^14^C studies, evidence from FACE experiments also suggest the existence of fine root populations with turnover of many years (Matamala *et al*., 2003; Korner *et al*., 2005; Keel *et al*., 2006; Lynch *et al*., 2013). Most of the isotopic studies deployed root screens to identify and measure newly grown roots with isotopic ages of 0–3 years to conclude that the old isotopic ages in standing mass must reflect longer lifetimes of the roots. However, it is clear that at least some of these studies did not necessarily separate tree roots from herbaceous or understory plants in the screens. While the ^14^C-age difference between standing biomass and screen roots in this study was small, the age distribution of known-age screen roots skewed younger (Fig. 6). The different age distributions can be explained if the screen roots contained a high percentage of ephemeral summer roots (with young ^14^C-ages) that contributed little to the standing-root biomass. Overall, our results add another piece of evidence that old isotopic ages of fine roots reflect mostly use of storage NSC and not slow root turnover (Solly *et al*., 2018).

## Conclusions

In the Alpine treeline ecotone we studied, young CO_2_ respired by the roots indicates that fresh NSC was transported and used belowground. At the same time, plant tissues also contained older NSC fixed on average years earlier, with increasing ages from needles to roots. Our mixing model suggests this vertical trend stems from higher influx of fresh NSC in the above-over the belowground tissues. The stored NSC was not intact and supported the growth of fine roots, which probably took place early in the season. The NSC aging towards the roots varied across elevations with the strongest aging in the bottom of the ecotone where a greater proportion of fresh NSC was used to support aboveground growth. At the top of the ecotone the aging was smaller, explained by the temperature limitation of aboveground growth that allowed a greater proportion of fresh NSC to flow to the roots. Minor species effects further indicated that environmental factors are more important for NSC allocation. We suggest that NSC ^14^C-ages provide a multi-year integrator of NSC allocation, with fine roots providing the most sensitive tissue for recording variations in belowground allocation.

## Acknowledgments

We thank Axel Steinhof and Heike Machts for processing and measuring ^14^C samples, Valérie F Schwab and Jose David Urquiza Muñoz for help in sample collection, Stephanie Strahl and Christin Leschik for NSC extractions, and Anette Enke for carbohydrates measurements. We are grateful to the “Stillberg” team at WSL/SNF to maintain the long-term afforestation experiment. Emily Solly acknowledges funding from the Swiss National Science Foundation (Ambizione grant PZ00P2_180030). Funding: The Max Planck Institute for Biogeochemistry and the European Research Council Horizon 2020 Research and Innovation Programme, grant agreement 695101 (14Constraint).

## Author contributions

BH, EFS, FH, and ST conceived the ideas and designed the study. BH, EFS, and FH collected the samples. BH measured respiration, IK performed NSC analysis, and EFS analyzed fine-root chronological ages. BH perfumed the data analysis and led the writing. All authors contributed to data interpretation and text editing.

## Data availability

The data that support the findings of this study are openly available in “figshare” at http://doi.org/10.6084/m9.figshare.24609192.v1

## Supporting information

**Table S1** Results of linear mixed-effects models testing effects on ^14^C ages.

## References

Andreu-Hayles L, Santos GM, Herrera-Ramírez DA, Martin-Fernández J, Ruiz-Carrascal D, Boza-Espinoza TE, Fuentes AF, J⊘rgensen PM. 2015. Matching Dendrochronological Dates with the Southern Hemisphere 14C Bomb Curve to Confirm Annual Tree Rings in Pseudolmedia rigida from Bolivia. Radiocarbon 57(1): 1–13.

Barbeito I, Dawes MA, Rixen C, Senn J, Bebi P. 2012. Factors driving mortality and growth at treeline: a 30-year experiment of 92⍰000 conifers. Ecology 93(2): 389–401.

Bates D, Mächler M, Bolker B, Walker S. 2015. Fitting Linear Mixed-Effects Models Using lme4. Journal of Statistical Software 67(1): 1–48.

Brunner I, Bakker MR, Bjork RG, Hirano Y, Lukac M, Aranda X, Borja I, Eldhuset TD, Helmisaari HS, Jourdan C, et al. 2013. Fine-root turnover rates of European forests revisited: an analysis of data from sequential coring and ingrowth cores. Plant and Soil 362(1-2): 357–372.

Carbone MS, Czimczik CI, Keenan TF, Murakami PF, Pederson N, Schaberg PG, Xu X, Richardson AD. 2013. Age, allocation and availability of nonstructural carbon in mature red maple trees. New Phytologist 200(4): 1145–1155.

Carbone MS, Still CJ, Ambrose AR, Dawson TE, Williams AP, Boot CM, Schaeffer SM, Schimel JP. 2011. Seasonal and episodic moisture controls on plant and microbial contributions to soil respiration. Oecologia 167(1): 265–278.

Cutler DF, Rudall P, Gasson P, Gale R. 1987. Root identification manual of trees and shrubs: a guide to the anatomy of roots of trees and shrubs hardy in Britain and Northern Europe. London, UK: Chapman and Hall.

D’Andrea E, Rezaie N, Battistelli A, Gavrichkova O, Kuhlmann I, Matteucci G, Moscatello S, Proietti S, Scartazza A, Trumbore S, et al. 2019. Winter’s bite: beech trees survive complete defoliation due to spring late-frost damage by mobilizing old C reserves. New Phytologist 224(2): 625–631.

Dawes MA, Philipson CD, Fonti P, Bebi P, Hattenschwiler S, Hagedorn F, Rixen C. 2015. Soil warming and CO2 enrichment induce biomass shifts in alpine tree line vegetation. Global Change Biology 21(5): 2005–2021.

Dietze MC, Sala A, Carbone MS, Czimczik CI, Mantooth JA, Richardson AD, Vargas R. 2014. Nonstructural carbon in woody plants. Annual Review of Plant Biology 65: 667–687.

Epron D, Bahn M, Derrien D, Lattanzi FA, Pumpanen J, Gessler A, Hogberg P, Maillard P, Dannoura M, Gerant D, et al. 2012. Pulse-labelling trees to study carbon allocation dynamics: a review of methods, current knowledge and future prospects. Tree Physiology 32(6): 776–798.

Farrar J, Hawes M, Jones D, Lindow S. 2003. How roots control the flux of carbon to the rhizosphere. Ecology 84(4): 827–837.

Ferrari A, Hagedorn F, Niklaus PA. 2016. Experimental soil warming and cooling alters the partitioning of recent assimilates: evidence from a 14C-labelling study at the alpine treeline. Oecologia 181(1): 25–37.

Gao D, Joseph J, Werner RA, Brunner I, Zürcher A, Hug C, Wang A, Zhao C, Bai E, Meusburger K, et al. 2021. Drought alters the carbon footprint of trees in soils—tracking the spatiotemporal fate of 13C-labelled assimilates in the soil of an old-growth pine forest. Global Change Biology 27(11): 2491–2506.

Gärtner H, Schweingruber F. 2013. Microscopic Preparation Techniques for Plant Stem Analysis. Remagen-Oberwinter, Germany: Verlag Dr. Kessel.

Gaudinski J, Trumbore S, Davidson E, Cook A, Markewitz D, Richter D. 2001. The age of fine-root carbon in three forests of the eastern United States measured by radiocarbon. Oecologia 129(3): 420–429.

Gaudinski JB, Torn M, Riley W, Dawson T, Joslin J, Majdi H. 2010. Measuring and modeling the spectrum of fine-root turnover times in three forests using isotopes, minirhizotrons, and the Radix model. Global Biogeochemical Cycles 24(3).

Guo DL, Li H, Mitchell RJ, Han WX, Hendricks JJ, Fahey TJ, Hendrick RL. 2008. Fine root heterogeneity by branch order: exploring the discrepancy in root turnover estimates between minirhizotron and carbon isotopic methods. New Phytologist 177(2): 443–456.

Hagedorn F, Martin M, Rixen C, Rusch S, Bebi P, Zürcher A, Siegwolf RTW, Wipf S, Escape C, Roy J, et al. 2010. Short-term responses of ecosystem carbon fluxes to experimental soil warming at the Swiss alpine treeline. Biogeochemistry 97(1): 7–19.

Hammer S, Levin I. 2017. Monthly mean atmospheric D14CO2 at Jungfraujoch and Schauinsland from 1986 to 2016. doi: 10.11588/data/10100

Handa IT, Körner C, Hättenschwiler S. 2005. A Test of the Treeline Carbon Limitation Hypothesis by in Situ CO2 enrichment and Defoliation. Ecology 86(5): 1288–1300.

Harkness DD, Harrison AF, Bacon PJ. 1986. The Temporal Distribution of ‘Bomb’ 14C in a Forest Soil. Radiocarbon 28(2A): 328–337.

Helmisaari H-S, Leppälammi-Kujansuu J, Sah S, Bryant C, Kleja D. 2015. Old carbon in young fine roots in boreal forests. Biogeochemistry 125(1): 37–46.

Herzog C, Steffen J, Graf Pannatier E, Hajdas I, Brunner I. 2014. Nine years of irrigation cause vegetation and fine root shifts in a water-limited pine forest. PLOS ONE 9(5): e96321.

Hilman B, Muhr J, Helm J, Kuhlmann I, Schulze ED, Trumbore S. 2021. The size and the age of the metabolically active carbon in tree roots. Plant, Cell & Environment 44(8): 2522–2535.

Hoch G, Körner C. 2009. Growth and carbon relations of tree line forming conifers at constant vs. variable low temperatures. Journal of Ecology 97(1): 57–66.

Hoch G, Körner C. 2012. Global patterns of mobile carbon stores in trees at the high-elevation tree line. Global Ecology and Biogeography 21(8): 861–871.

Högberg MN, Briones MJI, Keel SG, Metcalfe DB, Campbell C, Midwood AJ, Thornton B, Hurry V, Linder S, Näsholm T, et al. 2010. Quantification of effects of season and nitrogen supply on tree below-ground carbon transfer to ectomycorrhizal fungi and other soil organisms in a boreal pine forest. New Phytologist 187(2): 485–493.

Hua Q, Barbetti M, Rakowski AZ. 2013. Atmospheric Radiocarbon for the Period 1950–2010. Radiocarbon 55(4): 2059–2072.

Kagawa A, Sugimoto A, Maximov TC. 2006. Seasonal course of translocation, storage and remobilization of 13C pulse-labeled photoassimilate in naturally growing Larix gmelinii saplings. New Phytologist 171(4): 793–804.

Karst J, Gaster J, Wiley E, Landhausser SM. 2016. Stress differentially causes roots of tree seedlings to exude carbon. Tree Physiology 37(2): 154–164.

Keel SG, Siegwolf RT, Jaggi M, Korner C. 2007. Rapid mixing between old and new C pools in the canopy of mature forest trees. Plant, Cell & Environment 30(8): 963–972.

Keel SG, Siegwolf RT, Korner C. 2006. Canopy CO2 enrichment permits tracing the fate of recently assimilated carbon in a mature deciduous forest. New Phytologist 172(2): 319–329.

Körner C. 1998. A re-assessment of high elevation treeline positions and their explanation. Oecologia 115(4): 445–459.

Körner C 2003a. Alpine treelines. In: Alpine Plant Life: Functional Plant Ecology of High Mountain Ecosystems. Germany: Sprimger Berlin, Heidelberg, 77–100.

Körner C 2003b. Uptake and loss of carbon. In: Alpine Plant Life: Functional Plant Ecology of High Mountain Ecosystems. Germany: Sprimger Berlin, Heidelberg, 171–200.

Korner C, Asshoff R, Bignucolo O, Hattenschwiler S, Keel SG, Peláez-Riedl S, Pepin S, Siegwolf RT, Zotz G. 2005. Carbon flux and growth in mature deciduous forest trees exposed to elevated CO2. Science 309(5739): 1360–1362.

Kuznetsova A, Brockhoff PB, Christensen RHB. 2017. lmerTest Package: Tests in Linear Mixed Effects Models. Journal of Statistical Software 82(13): 1–26.

Lacointe A, Kajji A, Daudet FA, Archer P, Frossard JS. 1993. Mobilization of Carbon Reserves in Young Walnut Trees. Acta Botanica Gallica 140(4): 435–441.

Landhäusser SM, Chow PS, Dickman LT, Furze ME, Kuhlman I, Schmid S, Wiesenbauer J, Wild B, Gleixner G, Hartmann H, et al. 2018. Standardized protocols and procedures can precisely and accurately quantify non-structural carbohydrates. Tree Physiology 38(12): 1764–1778.

Leavitt SW, Danzer SR. 1993. Method for Batch Processing Small Wood Samples to Holocellulose for Stable-Carbon Isotope Analysis. Analytical Chemistry 65(1): 87–89.

Lenth RV 2021. Emmeans: estimated marginal means, aka least-squares means. R package version 1.5.4.

Litton CM, Raich JW, Ryan MG. 2007. Carbon allocation in forest ecosystems. Global Change Biology 13(10): 2089–2109.

Lynch DJ, Matamala R, Iversen CM, Norby RJ, Gonzalez-Meler MA. 2013. Stored carbon partly fuels fine-root respiration but is not used for production of new fine roots. New Phytologist 199(2): 420–430.

Marshall JD, Tarvainen L, Zhao P, Lim H, Wallin G, Nasholm T, Lundmark T, Linder S, Peichl M. 2023. Components explain, but do eddy fluxes constrain? Carbon budget of a nitrogen-fertilized boreal Scots pine forest. New Phytologist 239(6): 2166–2179.

Matamala R, Gonzalez-Meler MA, Jastrow JD, Norby RJ, Schlesinger WH. 2003. Impacts of fine root turnover on forest NPP and soil C sequestration potential. Science 302(5649): 1385–1387.

McCormack ML, Dickie IA, Eissenstat DM, Fahey TJ, Fernandez CW, Guo D, Helmisaari HS, Hobbie EA, Iversen CM, Jackson RB. 2015. Redefining fine roots improves understanding of below-ground contributions to terrestrial biosphere processes. New Phytologist 207(3): 505–518.

McNeely R. 1994. Long-term environmental monitoring of 14C levels in the Ottawa region. Environment International 20(5): 675–679.

Mrak T, Gričar J. 2016. Atlas of woody plant roots: morphology and anatomy with special emphasis on fine roots. Ljubljana, Slovenia: Slovenian Forestry Institute, The Silva Slovenica Publishing Centre.

Muhr J, Angert A, Negrón-Juárez RI, Muñoz WA, Kraemer G, Chambers JQ, Trumbore SE. 2013. Carbon dioxide emitted from live stems of tropical trees is several years old. Tree Physiology 33(7): 743–752.

Muhr J, Trumbore S, Higuchi N, Kunert N. 2018. Living on borrowed time - Amazonian trees use decade-old storage carbon to survive for months after complete stem girdling. New Phytologist 220(1): 111–120.

Peltier DMP, Lemoine J, Ebert C, Xu X, Ogle K, Richardson AD, Carbone MS. 2023. An incubation method to determine the age of available nonstructural carbon in woody plant tissues. Tree Physiology.

Prescott CE, Grayston SJ, Helmisaari HS, Kastovska E, Korner C, Lambers H, Meier IC, Millard P, Ostonen I. 2020. Surplus Carbon Drives Allocation and Plant-Soil Interactions. Trends in Ecology & Evolution 35(12): 1110–1118.

R Development Core Team. 2019. R: a language and environment for statistical computing. Vienna, Austria: R Foundation for Statistical Computing. URL https://www.R-project.org

Raessler M, Wissuwa B, Breul A, Unger W, Grimm T. 2010. Chromatographic analysis of major non-structural carbohydrates in several wood species – an analytical approach for higher accuracy of data. Analytical Methods 2(5): 532–538.

Richardson AD, Carbone MS, Keenan TF, Czimczik CI, Hollinger DY, Murakami P, Schaberg PG, Xu X. 2013. Seasonal dynamics and age of stemwood nonstructural carbohydrates in temperate forest trees. New Phytologist 197(3): 850–861.

Riley WJ, Gaudinski JB, Torn MS, Joslin JD, Hanson PJ. 2009. Fine-root mortality rates in a temperate forest: estimates using radiocarbon data and numerical modeling. New Phytologist 184(2): 387–398.

Rog I, Hilman B, Fox H, Yalin D, Qubaja R, Klein T. 2024. Increased belowground tree carbon allocation in a mature mixed forest in a dry versus a wet year. Global Change Biology 30(2): e17172.

Sah SP, Jungner H, Oinonen M, Kukkola M, Helmisaari HS. 2011. Does the age of fine root carbon indicate the age of fine roots in boreal forests?Biogeochemistry 104(1): 91–102.

Solly EF, Brunner I, Helmisaari HS, Herzog C, Leppalammi-Kujansuu J, Schoning I, Schrumpf M, Schweingruber FH, Trumbore SE, Hagedorn F. 2018. Unravelling the age of fine roots of temperate and boreal forests. Nature Communications 9(1): 3006.

Solly EF, Jaeger ACH, Barthel M, Werner RA, Zurcher A, Hagedorn F, Six J, Hartmann M. 2023. Water limitation intensity shifts carbon allocation dynamics in Scots pine mesocosms. Plant and Soil 490(1-2): 499–519.

Solly EF, Lindahl BD, Dawes MA, Peter M, Souza RC, Rixen C, Hagedorn F. 2017. Experimental soil warming shifts the fungal community composition at the alpine treeline. New Phytologist 215(2): 766–778.

Steinhof A, Altenburg M, Machts H. 2017. Sample Preparation at the Jena 14C Laboratory. Radiocarbon 59(3): 815–830.

Strand AE, Pritchard SG, McCormack ML, Davis MA, Oren R. 2008. Irreconcilable differences: fine-root life spans and soil carbon persistence. Science 319(5862): 456–458.

Streit K, Siegwolf RT, Hagedorn F, Schaub M, Buchmann N. 2014. Lack of photosynthetic or stomatal regulation after 9 years of elevated [CO2] and 4 years of soil warming in two conifer species at the alpine treeline. Plant, Cell & Environment 37(2): 315–326.

Tarkowski LP, Van den Ende W. 2015. Cold tolerance triggered by soluble sugars: a multifaceted countermeasure. Front Plant Sci 6: 203.

Trumbore S, Czimczik CI, Sierra CA, Muhr J, Xu X. 2015. Non-structural carbon dynamics and allocation relate to growth rate and leaf habit in California oaks. Tree Physiology 35(11): 1206–1222.

Trumbore S, Da Costa ES, Nepstad DC, Barbosa De Camargo P, Martinelli LA, Ray D, Restom T, Silver W. 2006. Dynamics of fine root carbon in Amazonian tropical ecosystems and the contribution of roots to soil respiration. Global Change Biology 12(2): 217–229.

Trumbore S, Sierra C, Pries CH 2016. Radiocarbon nomenclature, theory, models, and interpretation: measuring age, determining cycling rates, and tracing source pools. In: Schuur E, Druffel E, Trumbore S, eds. Radiocarbon and climate change. Switzerland: Sprimger, 45–82.

Vargas R, Trumbore SE, Allen MF. 2009. Evidence of old carbon used to grow new fine roots in a tropical forest. New Phytologist 182(3): 710–718.

